# Reliability of Static and Dynamic Network Metrics in the Resting-State: A MEG-beamformed Connectivity Analysis

**DOI:** 10.1101/358192

**Authors:** S. I. Dimitriadis, B. Routley, D. Linden, K.D. Singh

**Affiliations:** Cardiff University Brain Research Imaging Centre, School of Psychology, Cardiff University, Cardiff, United Kingdom; Neuroinformatics Group, Cardiff University Brain Research Imaging Centre, School of Psychology, Cardiff University, Cardiff, United Kingdom; Division of Psychological Medicine and Clinical Neurosciences, School of Medicine, Cardiff University, Cardiff, United Kingdom; School of Psychology, Cardiff University, Cardiff, United Kingdom; Neuroscience and Mental Health Research Institute, Cardiff University, Cardiff, United Kingdom; MRC Centre for Neuropsychiatric Genetics and Genomics, School of Medicine, Cardiff University, Cardiff, United Kingdom

**Keywords:** MEG, resting-state, time-varying network analysis, chronnectomics, functional connectivity microstates, symbolic analysis, reproducibility

## Abstract

The resting activity of the brain can be described by so-called intrinsic connectivity networks (ICNs), which consist of spatially and temporally distributed, but functionally connected, nodes. The coordinated activity of the resting state can be explored via magnetoencephalography (MEG) by studying frequency-dependent functional brain networks at the source level. Although many algorithms for the analysis of brain connectivity have been proposed, the reliability of network metrics derived from both static and dynamic functional connectivity is still unknown. This is a particular problem for studies of associations between ICN metrics and personality variables or other traits, and for studies of differences between patient and control groups, which both depend critically on the reliability of the metrics used. A detailed investigation of the reliability of metrics derived from resting-state MEG repeat scans is therefore a prerequisite for the development of connectomic biomarkers.

Here, we first estimated both static (SFC) and dynamic functional connectivity (DFC) after beamforming source reconstruction using the imaginary part of the phase locking index (iPLV) and the correlation of the amplitude envelope (CorEnv). Using our approach, functional network microstates (FCμstates) were derived from the DFC and chronnectomics were computed from the evolution of FCμstates across experimental time. In both temporal scales, the reliability of network metrics (SFC), the FCμstates and the related chronnectomics were evaluated for every frequency band.

Chronnectomic parameters and FCμstates were generally more reliable than node-wise static network metrics. CorEnv-based network metrics were more reproducible at the static approach. This analysis encourages the analysis of MEG resting-state via DFC.

## 1. Introduction

The coordination of spontaneous activity can be characterized with functional connectivity (FC), which refers to statistical dependencies between the activity of distinct brain areas (Pereda et al., 2005) and has been linked to the efficiency of an individual’s brain functioning (Baldassarre et al., 2012; Yamashita et al., 2015).

A functional connectivity graph (FCG) can be constructed by estimating the statistical dependencies between the brain activity of all the areas in a pair-wise fashion. An FCG represents statistical or causal relationships measured as cross-correlations, coherence, or information flow.

Neuroscientists first examined resting-state FC with functional magnetic resonance imaging (fMRI) by correlating blood oxygenation level-dependent (BOLD) signals (Biswal et al., 1995, 2011, 2012; van den Heuvel et al., 2009). After twenty years of using fMRI as a dominant neuroimaging tool, the community has succeeded in mapping brain areas to specific brain functions, creating an anatomical-functional atlas (Bandettini, 2012). Although fMRI is of high interest and a key modality to explore human brain function, ultra-slow activity described via BOLD signals is only an indirect measure of brain activity (Logothetis, 2008).

In the last few years, greater attention has been given to explore FC via electro-magneto-encephalography. Even though the spatial resolution of magnetoencephalography (MEG) is lower when compared to fMRI, MEG can capture the multiplexity of human brain activity by providing insight into the spectro-temporo-spatial dynamics of human brain activity. MEG-based FC provides us with a direct measure of neuromagnetic activity with a high temporal resolution.

Resting-state networks (RSNs) have been successfully extracted with MEG over the past few years using source-space FC (Brookes et al., 2011a, b; de Pasquale et al., 2010; Hall et al., 2013; Hipp et al., 2012; Luckhoo et al., 2012; Wens et al., 2014). Moreover, resting-state MEG FC has been proven to detect abnormal brain functioning in a variety of diseases, including Alzheimer’s disease (Engels et al., 2015; López et al., 2014,2017), multiple sclerosis (Tewarie et al., 2015), in schizophrenia (Bowyer et al., 2015), in dyslexia (Dimitriadis et al., 2013b,2015c), in mild cognitive impairment (Dimitriadis et al., 2015b) and in mild traumatic brain injury (Dunkley et al., 2015; Dimitriadis et al., 2015c, Antonanakis et al., 2016, 2017).

Several studies have thus captured alterations of MEG parameters in the resting state in order to estimate FC in disease groups compared to controls. However, FC estimates at resting-state could be affected by subject’s cognitive, emotional state and other scanning-related systematic differences. For that reason, it is unclear up to which level FC estimates are repeatable for an individual. Moreover, in large studies of hundreds of participants, there is a significant cost, both in financial resources and time, to scan all the subjects two or more times. To establish MEG as a clinically reliable neuroimaging tool that can distinguish disease from healthy populations, the reliability of FC patterns should be explored from repeat scans. Up to date, only a few studies accessed the test-retest reliability of MEG/electroencephalography (EEG) FC (Hardmeier et al., 2014; Jin et al., 2011; Garces et al., 2016) while only one study has quantified the test-retest reliability of FC estimates in the source-space MEG (Garces et al., 2016). Colclough et al., (2016) attempted to report the reliability of every edge-weighted connections with a high number of connectivity estimators but using a split-half strategy from a large pool of subjects. Practically, the results cannot be adopted as reliability of static network metrics since the analysis involved single MEG scan recordings. However, no study has ever explored the reliability of both static and dynamic networks at the source space in MEG.

In the present study, we investigated the test-retest reliability of both static and dynamic FC measures derived from MEG resting-state data. For that purpose, we computed whole-brain FC for 40 subjects who were scanned twice with a 1-week test-retest interval. For each subject and session, MEG-beamformed source activity was estimated and FC was computed between 90 brain areas. FC was estimated with the imaginary part of the phase locking index (iPLV) and the correlation of the amplitude envelope (CorEnv) in both static (SFC) and dynamic models (DFC) by adopting a sliding window approach (De Pasquale et al., 2010; Dimitriadis *et al.*, 2010a,2012a,2013a,2015a,2016a,2017). Afterwards, statistical and topological filtering schemes were applied to both SFC and DFC to reveal the true topology (Dimitriadis et al., 2017). For the SFC approach, we estimated well-known network metrics in a node-wise fashion and the reliability was accessed via correlation values between the two measurements and across the cohort. Graph-based reliability was assessed with a novel graph diffusion distance metric.

For the DFC approach, node-wise network metrics were estimated across experimental time. To explore spatio-temporally the derived network activity, we first designed a codebook of prototypical network microstates and then assigned each of the instantaneous connectivity patterns to the most similar code symbol (e.g. functional connectivity graph – FCG) (Dimitriadis *et al.*, 2010a,2012a,2013a, b,2015a,2016a, b,2017). A codebook is a set of prototypical functional connectivity graphs (FCGs). In this way, we derive a unique symbolic time series from each individual where each symbol corresponds to one of the predefined prototypical functional connectivity microstates (FCμstates). The evolution of these symbol-patterns encapsulates significant state transitions. Furthermore, the evolution of these FCμstates can be seen as a first order Markovian Chain (MC) that can be modelled representing an individualized state transition model of resting-state FCμstates. Fractional occupancy of each FCμstate, transition rates of FCμstates and MC models are the key features to explore the reliability of chronnectome in MEG source space. The group-consistency of subject-specific FCμstates was further explored. The whole analysis of dynamic functional connectivity graphs and the definition of FCμstates have been described in previous paper (Dimitriadis et al., 2013a).

Many techniques have already been proposed to summarize brain activity into short-lived transient brain states using the spectrum of neuromagnetic recordings (Vidaurre et al., 2016) and also the band-limited amplitude envelopes of source reconstructed MEG data (Baker et al., 2014; O’Neill et al., 2015). In detail, Vidaurre et al.,(2016) proposed a combination of multivariate autoregressive model with hidden markovian modelling (MAR-HMM) in order to model the temporal, spectral and spatial properties of MEG reconstructed activity into very short-lived brain states. Similarly, Baker et al., (2014) modelled resting-state source-reconstructed MEG activity with HMM into distinct spatio-temporal activation profiles called brain states. These brain states were linked to well-known anatomical brain areas. O’Neill et al.,(2015) mined MEG source activity from two tasks, a self-paced motor and a Sternberg working memory task. He used a sliding window canonical correlation analysis (CCA) to estimate the functional connectivity at each time-window and a k-means clustering to detect repeatable spatial patterns of connectivity that form transiently synchronising sub-networks (TSNs) or functional connectivity microstates. Here, we must underline the distinction of summarizing brain activity using the raw time series (ROIs × sliding windows; Vidaurre et al., 2016; Baker et al., 2014) which is a 2D matrix and the dynamic functional brain networks (ROIs × ROIs × sliding windows) which is a 3D matrix (O’Neill et al.,2015). Currently the mapping and the relationship between raw activity and brain connectivity and also the relationship of microstates (raw activity) with functional connectivity microstates (dynamic graphs; Allen et al., 2012; Dimitriadis *et al.*, 2013a, b,2015a,2016a, b,2017) is still unknown. Further research is need to explore their mapping at resting-state and during tasks.

The proposed methodological scheme entails two distinct ways of analyzing dynamic functional connectivity patterns. These patterns are representative brain network topologies across subjects and brain rhythms and are directly linked to a brain state. The very first approaches in fMRI constitutes novel contributions to an emerging neuroimaging field called chronnectomics (Allen et al., 2012; Calhoun *et al.*, 2014). Previously, we reported the notion of FCμstates (Dimitriadis et al., 2013a) and the developmental trends in cognition (Dimitriadis et al., 2015a) using electroencephalographic recordings. The concept of chronnectome is the incorporation of a dynamic view of functional brain connectivity networks and the evolution of revisiting distinct spatio-temporal brain states (functional connectivity microstates (FCμstates). To the best of our knowledge, this study constitutes the first attempt to assess the test-retest reliability of Dynamic Functional Connectivity at the MEG source level.

Despite growing enthusiasm in the neuroscience community about the potential contribution of neuroimaging and especially brain networks in the designing of connectomic biomarkers for various brain diseases/disorders, many challenges remain open (Stam et al.,2014). At first level, it is more than significant to explore how reliable are network metrics at both temporal scales (static and dynamic) by analysing a group of control subjects with repeat scans (e.g. diffusion MRI: Dimitriadis et al., 2017d). Here, we assess evidence of the reliability of neuromagnetic (MEG) based functional connectomics to lead to potential clinically meaningful biomarker identification in target populations through the lens of the criteria used to evaluate clinical tests.

## 2. Materials and Methods

### 2.1 Subjects

40 healthy subjects (age *22¨85*±*3.74* years, 15 women and 25 men) underwent two resting-state MEG sessions (eyes open) with a 1-week test-retest interval. For each participant, scans were scheduled at the same day of the week and same time of the day. The duration of MEG resting-state was 5 mins for every participant. The study was approved by the Ethics Committee of the School of Psychology at Cardiff University, and participants provided informed and written consent.

### 2.2 MEG-MRI Recordings

Whole-head MEG recordings were made using a 275-channel CTF radial gradiometer system. An additional 29 reference channels were recorded for noise cancellation purposes and the primary sensors were analysed as synthetic third-order gradiometers (Vrba and Robinson, 2001). Two or three of the 275 channels were turned off due to excessive sensor noise (depending on time of acquisition). Subjects were seated upright in the magnetically shielded room. To achieve MRI/MEG co-registration, fiduciary markers were placed at fixed distances from three anatomical landmarks identifiable in the subject’s anatomical MRI, and their locations were verified afterwards using high-resolution digital photographs. Head localisation was performed before and after each recording, and a trigger was sent to the acquisition computer at relevant stimulus events.

All datasets were either acquired at or down-sampled to 600 Hz, and filtered with a 1-Hz high-pass and a 200-Hz lowpass filter. The data were first whitened and reduced in dimensionality using principal component analysis with a threshold set to 95% of the total variance (Delorme and Makeig, 2004). The statistical values of kurtosis, Rényi entropy and skewness of each independent component were used to eliminate ocular and cardiac artifacts. Specifically, a component was deemed artifactual if more than 20% of its values after normalization to zero-mean and unit-variance were outside the range of [−2, +2] (Delorme and Makeig, 2004; Escudero et al., 2011; Antonakakis et al., 2016). The artifact-free multichannel MEG resting-state recordings were then entered in the beamforming analysis (see next section).

Subjects further underwent an MRI session in which a 3T GE scanner with an 8-channel receive-only head RF coil T1- weighted 1-mm anatomical scan was acquired, using an inversion recovery spoiled gradient echo acquisition.

### 2.3 Beamforming

An atlas-based beamformer approach was adopted to project MEG data from the sensor level to source space independently for each brain rhythm. The frequency bands studied were: δ (0.5–4 Hz), θ (4–8 Hz), α_1_ (8–10 Hz), α_2_ (10–13 Hz), β_1_ (13–20 Hz), β_2_ (20–30 Hz), γ_1_ (30–45 Hz), γ_2_ (55–90 Hz). First, the coregistered MRI was spatially normalized to a template MRI using SPM8 (Weiskopf et al., 2011). The AAL atlas was used to anatomically label the voxels, for each participant and session, in this template space (Tzourio-Mazoyer N, et al., 2002). The 90 cortical regions of interest (ROIs) were used for further analysis, as is common in recent studies (Hillebrand et al., 2016, Hunt et al., 2016). Next, neuronal activity in the atlas-labelled voxels was reconstructed using the LCMV source localization algorithm as implemented in Fieldtrip (Oostenveld et al., 2012).

The beamformer sequentially reconstructs the activity for each voxel in a predefined grid covering the entire brain (spacing 6 mm) by weighting the contribution of each MEG sensor to a voxel’s time series - a procedure that creates the spatial filters that can then project sensor activity to the cortical activity. Each ROI in the atlas contains many voxels, and the number of voxels per ROI differs. To obtain a single representative time series for every ROI, we defined a functional-centroid ROI representative by functionally interpolating activity from the voxel time series, within each ROI, in a weighted fashion. Specifically, we estimated a functional connectivity map between every pair of source time series within each of the AALs ROIs (eq.1) using correlation (eq.2). We then estimated the connectivity strength of each voxel within the ROI by summing its connectivity values to other voxels within the same ROI (eq.3) and finally we normalized each strength by the sum of strengths (eq.4) to estimate a set of weights within the ROI that sum to a value of 1. Finally, we multiplied each voxel time series with their respective weights and we summed across them in order to get a representative time series for each ROI (eq.5). The whole procedure was applied independently to every quasi-stable temporal segment derived by the settings of temporal window and stepping criterion.

The following equations 1–5 demonstrated the steps for this functional interpolation.

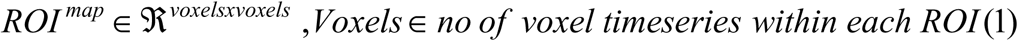

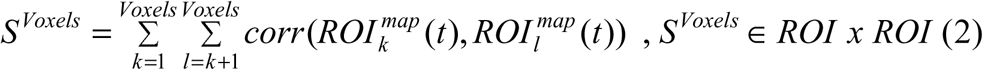

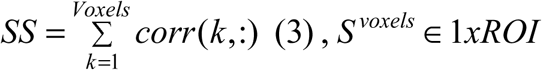

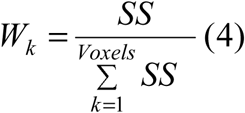

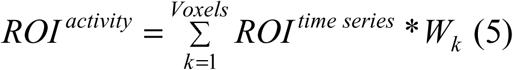

The outline of the methodology is described in Fig.1. An exemplar of the representative ban filtered ROI based time series is given in Fig.1. Fig.2 illustrates the preprocessing steps describ equations 1–5.

**Figure 1.**
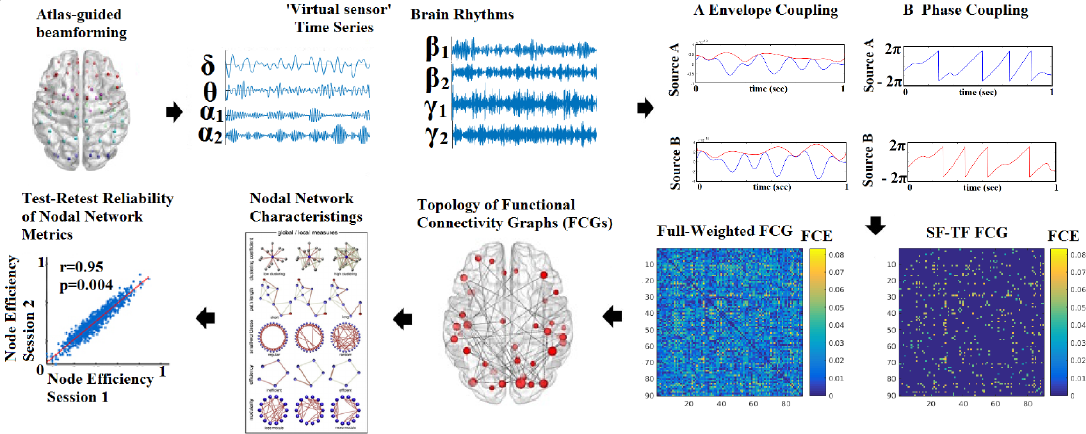
Outline of the methodology for accessing the reliability of network metrics derived from functional connectivity graphs (FCGs). (SF-Statistical Filtering, TF-Topological Filtering, FCE-Function Connectivity Estimator).

**Figure 2.**
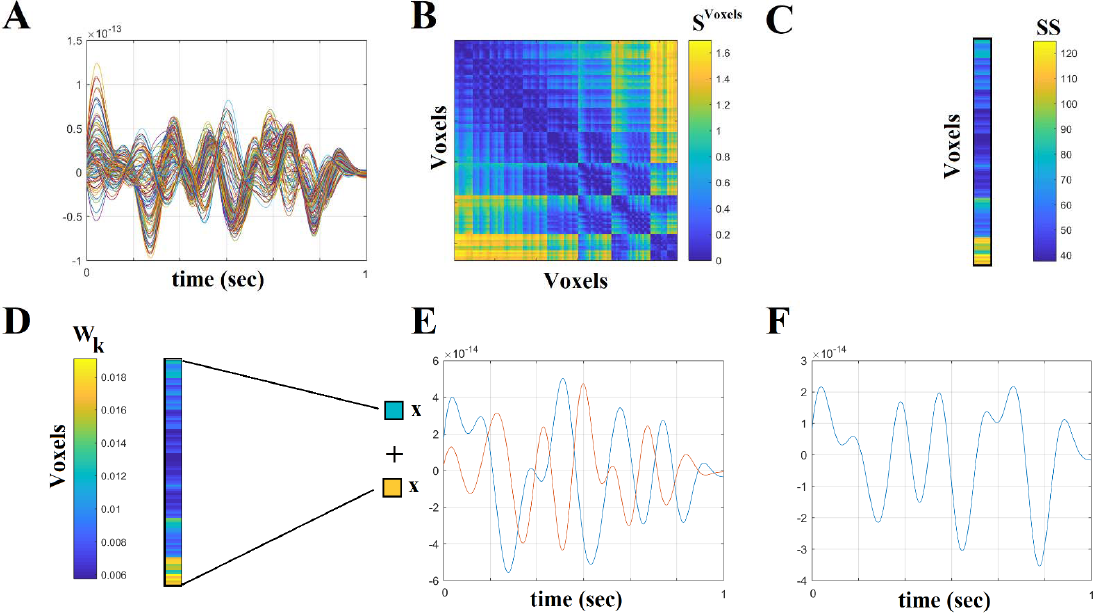
Step-by-step construction of the representative virtual sensor time series for each ROI. **A) Plot of 108 voxel time series from left precentral gyrus.** **B) Distance correlation matrix** *S ^Voxels^*derived by the pair-wise estimation of the 108 voxel time serie **C) Summation of the columns of***^SVoxels^*produced the vector SS **D) Normalization of vector SS that further produces W**_k_ where its sum equals to 1. **E) Multiplying every voxel time series with the related weight from the W_k_. In this example, w demonstrated this multiplication for the first and last voxel time series.** **F) The reproduced voxel time series for left precentral gyrus ((ROI** ^activity^). **after summing the weighted versions of every voxel time series from E.**

**Statistical and topologically filtering of the FCGs will be described in sections 2.7 and 2.8, correspondingly. One can understand how from a full-weighted FCG, a more sparse version is deriv via the statistical and topological filtering approaches.**

### 2.3 Functional Connectivity

Here, functional connectivity was examined among the following 8 brain rhythms of the typical sub-bands of electrophysiological neural signals {δ, θ, α_1_, α_2,_ β_1_, β_2_, γ_1_, γ_2_}, defined respectively within the ranges {0.5–4 Hz; 4–8 Hz; 8–10Hz; 10–13Hz; 13–20 Hz; 20–30 Hz; 30–45Hz; 55–90 Hz}. For both static and dynamic approach, we used two estimators: the correlation of the amplitude envelope (CorEnv) and the imaginary part of the phase locking value (iPLV).

### 2.4 Intra-Frequency Connectivity Estimators

Among the available connectivity estimators, we adopted one based on the imaginary part of phase-locking value (iPLV) (Lachaux *et al.*, 1999) and adjusted properly so as to extract time-resolved profiles of intra-frequency coupling from MEG multichannel recordings at resting state. The original PLV is defined as follows:

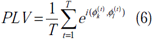

where k, l denote a pair of MEG sources and the imaginary part of PLV is equal to:

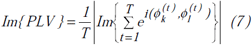

The imaginary part of PLV (iPLV) investigates intra-frequency interactions without putative contributions from volume conductance. In general, the iPLV is mainly sensitive to non-zero-phase lags and for that reason is resistant to instantaneous self-interactions from volume conductance (Nolte, 2004). In contrast, it could be sensitive to phase changes that not necessarily imply a PLV oriented coupling.

Correlation of the Envelope coupling (CorEnv) is based upon correlation between the oscillatory envelopes of two band limited sources (Brookes et al., 2012). See Fig.1 for a schematic diagram of phase and envelope based connectivity analyses based upon neural oscillations. Correlation of the Envelope coupling (CorEnv) is based upon correlation between the oscillatory envelopes of two band limited sources (A) while phase coupling searches for a constant phase lag between signals, in the example a difference of π (B). The time series for the estimation of CorEnv were orthogonalized between each other using the bivariate version of this correction for signal leakage effects (Colclough et al., 2016).

### 2.5 Static Functional Connectivity Analysis

Using both connectivity estimators, we estimated the fully-weighted (90×90) anatomical oriented FCG, one for each subject, recording session and frequency band. To construct the static FCG (SFCG), we incorporated in the analysis the whole 5 minutes of the recording session.

### 2.6 Dynamic iPLV estimates: the Time-Varying Integrated iPLV graph (^TVI^iPLV graph)

The goal of the analytic procedures described in this section was to understand the repertoire of phase-to-phase interactions and their temporal evolution, while taking into account the quasi-instantaneous spatiotemporal distribution of iPLV estimates. This was achieved by computing one set of iPLV estimates within each of a series of sliding overlapping windows spanning the entire 5-min continuous MEG recording for eyes-open condition. The width of the temporal window and the stepping criterion were optimized for each frequency band separately using as objective criterion the reliability of transition dynamics between scan session 1 and 2 for each brain rhythm (Dimitriadis *et al.*, 2013a; see section 2.13.1 and 2.15). The centre of the stepping window moved forwards every frequency-dependent (see section 2.15 and 3.1) for the optimization of the parameters) for every intra-frequency interactions and a new functional brain network is re-estimated between every pair of ‘swifting’ temporal segments of MEG activity, from two sources, leading to a “quasi-stable in time” static iPLV graph. In this manner, a series of 598 (for δ) to 2140 (for γ_2_) iPLV graph estimates were computed for each frequency (8 within frequency), for each participant and for both repeat scans.

For each subject, a 4D tensor (frequencies bands (8) × slides (598 to 2140) × sources (90) × sources (90); see section 2.15 and 3.1) was created for each condition integrating subject-specific spatio-temporal phase interactions (Fig.3.A).

**Fig. 3.**
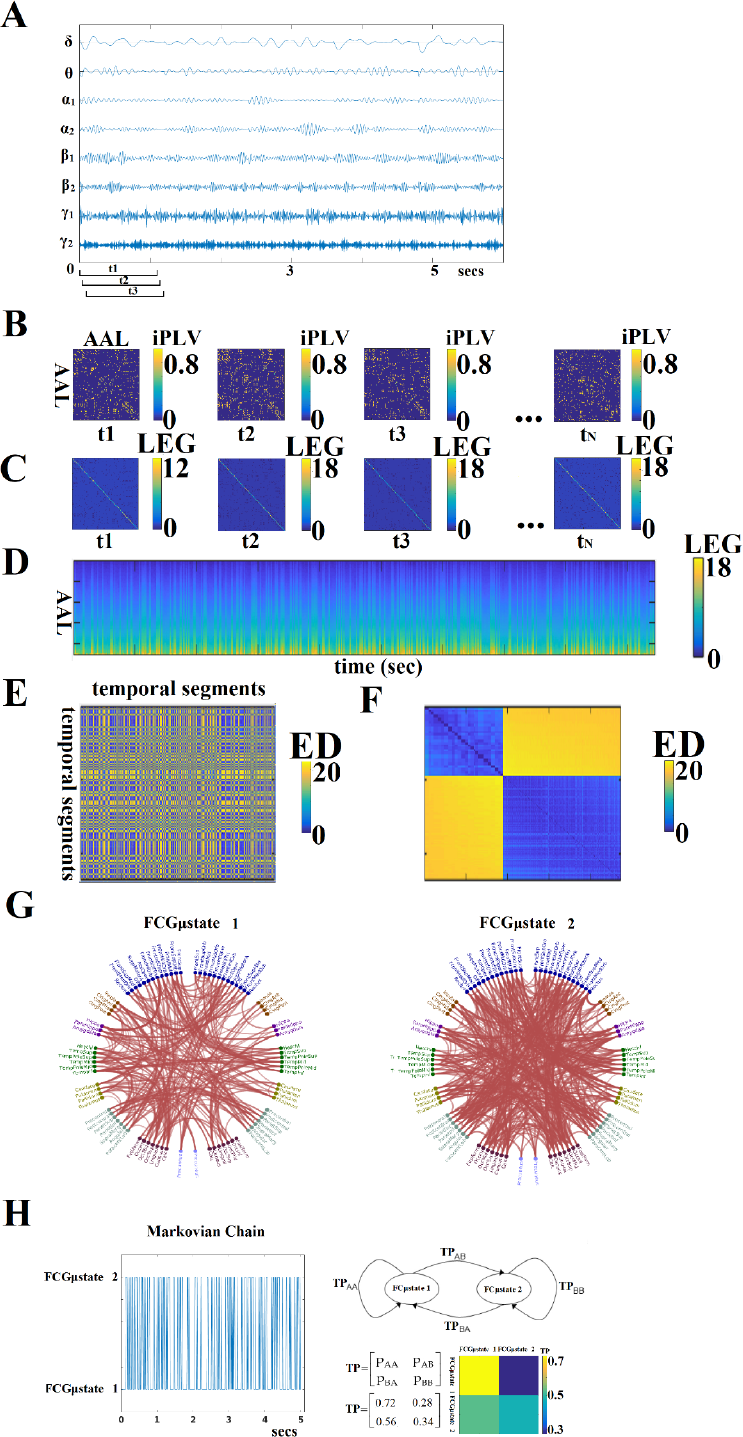
From dynamic functional connectivity graphs (DFCG) to FCμstates. A) A characteristic bandpass filtered time series for each of the studying frequency band is given from a ROI. B) Topologies of snapshots of DFCG from the three first temporal segments from δ band of subject 1 in order to make clear the estimation of FCG in a dynamic fashion. The first three brain networks refer to the first three temporal segments demonstrated in A). These functional brain networks were statistically and topologically filtered as described in sections 2.7 and 2.8. The t_N_ refers to the last temporal segment of the DFCG. C) Laplacian matrices for a few snapshots of DFCG D) The dynamic evolution of the eigenvalues of the laplacian matrices for each frequency band. An example for δ frequency band. E) Euclidean Distance matrix of the laplacian eigenvalues between every pair of temporal segments F) Reordering the correlation matrix in E to enhance the visualization of the two clusters – Fcμstates illustrated in G G) The prototypical Fcμstates in a circular visualization H) The outcome of this procedure is a symbolic time series that can be seen as a Markovian Chain that expresses the evolution of FCμstates across experimental time. The transition probabilities (TP) of this Markovian Chain based is illustrated in the 2 × 2 colored figure. One can understand that human brain demonstrates a preferred transition from FCμstates^2^ to FCμstates1 compared to the opposite direction (see 2D colormap). The chronnectomics were derived from this symbolic time series. (ED:Euclidean Distance, LEG:Laplacian EiGenvalues, ED:Euclideand Distance, AAL:Automated Anatomical Labeling LEG:Laplacian EiGenvalues)

### 2.7 Surrogate Data Analysis of *iPLV/CorEnv* Estimates – Statistical Filtering of Brain Networks

To identify significant *iPLV/CorEnv* -interactions which were estimated for every pair of frequencies within and between all 90 sources, and at each successive sliding window (i.e. temporal segment), we employed a surrogate data analysis. Accordingly, we could determine (a) if a given *iPLV/CorEnv* value differed from what would be expected by chance alone, and (b) if a non–zero *iPLV/CorEnv* corresponded to non–spurious coupling.

For every temporal segment, sensor–pair and frequency, we tested the null hypothesis H_0_: “the observed *iPLV/CorEnv* value comes from the same distribution as the distribution of surrogate *iPLV/CorEnv* -values”. One thousand surrogate time–series were generated by cutting at a single point at a random location the original time series and exchanging the two resulting time courses (Aru et al., 2015). We restricted the range of the selected cutting point in a temporal window of width to 10 sec in the middle of the recording session (between 25 – 35 sec). This surrogate scheme was applied to the original whole time series and not to the signal–segment at every slide. Repeating this procedure leads to a set of surrogates with a minimal distortion of the original phase dynamics, while the non–stationarity of the brain activity is less destroyed compared to shuffling the time series or cutting and rebuilding it in more than one time points.

This procedure ensures that the real and surrogate indices both have the same statistical properties. For each data set, the surrogate *iPLV/CorEnv (^s^iPLV/^s^CorEnv*) was then computed. We then determined a one–sided p–value for each *iPLV/CorEnv* value that corresponded to the likelihood that the observed value could belong to the surrogate distribution. This was done by directly estimating the proportion of “surrogate” *^s^iPLV/^s^*CorEnv that were higher than the observed *iPLV/CorEnv*–level (a very low value revealed that it could not have appeared from processes with no iPLV coupling).

At a second level, we applied the FDR method (Benjamini, Hochberg, 1995) to control for mu comparisons within each snapshot of the dynamic graph (FCG β a 90 × 90 matrix with tabulated p-values) with the expected fraction of false positives set to 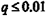. Finally, for each subject the res TV*iPLV/*^TV^*CorEnv* profiles constituted of two 3D arrays of size [598 to 2140 for δ to γ_2_ (time windows (sources) × 90 (sources)] with a value of 0 indicated a non-significant *iPLV/CorEnv* value.

The aforementioned statistical filtering approach was applied independently for each frequ band, session, subject and connectivity estimator for both static and dynamic functional connec graphs.

### 2.8 A data-driven Topological Filtering Scheme based on Orthogonal Minimal Spanning Trees (OMSTs)

As well as the statistical filtering approach, it is important to adopt a data-driven topolo filtering approach in order to reveal the backbone of the network topology over the increme information flow.

Recently, it was proved that MST is an unbiased method that yields reliable network metrics (Tewarie *et al.*, 2015). In this study, we adopt a variant of this topological filtering scheme called orthogonal mi spanning trees (OMST), which leads to a better sampling of brain networks, preserving the advanta MST, that connects the whole network with minimum cost without introducing cycles and wi differentiated strong from weak connections compared to the absolute threshold or the density thre (Dimitriadis et al., 2017a, b). MST is too sparse to capture the ‘true’ network and for that reason leading to the selection of N-1 connections where N denotes the number of nodes. We introduced OMST w samples the weights of a brain network via the notion of MST and under the optimization of the g information flow under the constraint of the total Cost of preserving the functional connec (Dimitriadis et al., 2017).

Our criterion for topologically filtering a given brain network is based on the maximum value of the following quality formula:

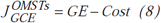

We applied the data-driven topological filtering scheme based on OMST at every static and quasi-instantaneous FCG from the dynamic DFCG. After statistical and topological filtering approaches applied to both SFCG and the DFCG, we estimated network metrics at the node/source level.

Fig.1 demonstrates an example of a full-weighted FCG after applying both statistical and topological filtering approach. Our algorithm was validated over all the existing thresholding schemes with a large EEG dataset over brain fingerprinting and with a multi-scan fMRI dataset over reliability of nodal network metrics (Dimitriadis et al., 2017a). Additionally, we demonstrated the importance of a data-driven topological filtering technique in functional neuroimaging by using OMST in a multi-group study with MEG resting-state recordings (Dimitriadis et al., 2017b). The MATLAB code of the OMST method and also of the majority of existing filtering methods can be downloaded from the: https://github.com/stdimitr/topological_filtering_networks & researchgate https://www.researchgate.net/profile/Stavros_Dimitriadis

### 2.9 Graph Diffusion Distance Metric for Brain Networks

In order to assess group and scan sessions differences in the topologically filtered FCG at the single-case level, we computed the Graph Diffusion Distance as a distance metric (Fouss et al., 2012; Hammond et al., 2013) from the OMST-derived final Functional Connectivity Graphs (FCG). The graph laplacian operator of each subject-specific FCG was defined as L = D – FCG, where D is a diagonal degree matrix related to FCG. This method entails modeling hypothetical patterns of information flow among sources based on each observed (static) SFCG. The diffusion process on the person-specific FCG was allowed for a set time t; the quantity that underwent diffusion at each time point is represented by the time-varying vector 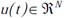. Thus, for a pair of sources i and j, the quantity FCG_ij_ (u_i_(t) – u_j_(t)) represents the hypothetical flow of information from i to j via the edges that connect them (both directly and indirectly). Summing all these hypothetical interactions for each sensor leads to 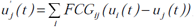, which can be written as:

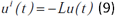

where L is the graph laplacian of FCG. At time t = 0 Equation 9 has the analytic solution: 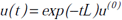. Here exp(-tL) is a N × N matrix function of t, known as Laplacian exponential diffusion kernel (Fouss et al., 2012), and u^(0)^ = e_j_, where 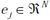is the unit vector with all zeros except in the j^th^ component. Running the diffusion process through time t produced the diffusion pattern exp(-tL)ejwhich corresponds to the jth column of exp(-tL).

Next, a metric of dissimilarity between every possible pair of person-specific diffusion-kernelized FCGs (FCG_1_, FCG_2)_ was computed in the form of the graph diffusion distance d_gdd_(t). The higher the value of d_gdd_(t) between two graphs, the more distinct is their network topology as well as the corresponding, hypothetical information flow. The columns of the Laplacian exponential kernels, exp(-tL1) and exp(-tL2), describe distinct diffusion patterns, centered at two corresponding sources within each FCG. The d_gdd_(t) function is searching for a diffusion time t that maximizes the Frobenius norm of the sum of squared differences between these patterns, summed over all sources, and is computed as:

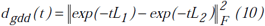

where 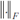is the Frobenius norm.

Given the spectral decomposition L=VΛV, the laplacian exponential can be estimated via

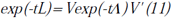

where for Λ, exp(-tΛ) is diagonal to the ith entry given by 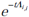. We computed d_gdd_(FCG_1_, FCG_2_) by first diagonalizing L1 and L2 and then applying equations (9) and (10) to estimate d_gdd_(t) for each time point t of the diffusion process. In this manner, a single dissimilarity value was computed for each pair of participants based on their individual characteristic FCGs. For further details see (Hammond et al., 2013). The GDD metric can be downloaded from:

https://github.com/stdimitr/multi-group-analysis-OMST-GDD

### 2.10 Static Network Metrics

After applying the statistical and topological filtering approach, we estimated the global effic for each node in static approach. The static approach leads to 90 (sources) values for each net metric, frequency band and session per subject. We adopted complementary features that measur importance of each node in segregation, integration and the information flow within a weig functional brain network (Dimitriadis *et al.*, 2010a, b,2013a, b,2015a). In this study, we estimated basic network metrics, the global and local efficiency, the strength of each node and the mean passage time based on random walks (Goni et al., 2013).

**Network global efficiency (GE)** reflects the overall efficiency of parallel information transfer w the entire set of 90 sources and was estimated as the average sourcespecific GE value over all so using the following formula (Latora and Marchiori, 2001): Where d denotes the shortest path length from i to j.

**Local efficiency** (LE) indicates how well the subgraphs exchange information when a part node is eliminated (Achard and Bullmore, 2007). Specifically, each node is assigned the shortest length within its subgraph where corresponds to the total number of spatial (first level neighbors) neighbors of the -th while d denotes shortest path length.

The **strength** is equal to the total sum of the weights of the connections of each node.

As a fourth candidate network metric, we adopted the mean first passage time (MFPT). Star random walk process on a brain network, an analytic expression can give the probability that a s particle departing from a node*i* arrives at node*j* for the first time within exactly*L* steps (Wang Pei,2008). This criterion can be applied for each MEG source pair by setting*L*to their shortest-path-length. We denote with 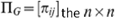the*n×n*symmetric matrix containing, for each pair of nodes probability of a single particle going from node *i* to node *j* via the shortest path. Each entry*π^i,j^*can be computed as

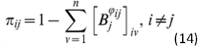

where matrix 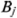is the transition matrix *P* introduced above, but with all zeros in the *j*-th colum, i.e. with *j* acting as an absorbing state (Wang and Pei, 2008). Evaluating shortest-path-lengths en that 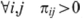. By considering one particle here, the average shortest-path probability of a gra defined as

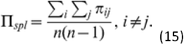

The derived 2D matrix based on nodal NMTS of GE, LE, MFPT and strength will be modelled the proposed method that is described in the following section.

### 2.11 Modelling of Dynamic Functional Connectivity Graphs (DFCG) as a 3D Tensor

This subsection serves as a brief introduction to our symbolization scheme, presented in gr details elsewhere (Dimitriadis et al., 2012a, b,2013a, b). The dynamic functional connectivity pattern be modeled as prototypical functional connectivity microstates (FCμstates). In a recent study demonstrated a better modeling of dynamic functional connectivity graphs (DFCG) based on a v quantization approach (Dimitriadis et al., 2013). In our previous work (Dimitriadis et al., 2013a, b,2015a), we used the neural-gas algorithm (Martinetz et al., 1993) to learn the 2D matrix (vectorized version matrix × time) leading to a codebook of *k* prototypical functional connectivity states (i.e. FCμstates) algorithm is an artificial neural network model, which converges efficiently to a small number codebook vectors, using a stochastic gradient descent procedure with a soft-max adaptation rule minimizes the average distortion error (Martinetz et al., 1993). In a recent study, we adopted negative matrix factorization (NNMF) as an appropriate learning algorithm of the 2D vectorized vers a dynamic functional brain network (Marimpis et al., 2016).

In our previous study where we first demonstrated how to model dynamic functional connectivity graph (dFCG) (Dimitriadis et al., 2013a), we vectorised the upper triangular of each of the quasi-static FCGs building a 2D matrix where the 1^st^ dimension is the number of temporal segments and the 2^nd^ the vectorised version of a static FCG. The final outcome of this approach is to define the so-called functional connectivity microstates (FCμstates). In a next study, we moved one step further by estimating node-wise global efficiency as the best descriptor to characterize the brain activity. The final outcome of the modelling using the same methodology of neural-gas algorithm was task-based network microstates (Dimitriadis et al., 2015a). Here, the vectorised version of a 90×90 FCG produces a long vector of 4005 values while the number of temporal segments ranged from 598 to 2140 which caused the so-called curse of dimensionality where the number of number of the temporal segment over which the modelling will learn the brain states is much smaller compared to the vectorised snapshot of FCG. Simultaneously, the vectorised notion of a brain network didn’t maintain the inherent format of a functional brain network which is a 2D matrix, a tensor.

The outline of this procedure is illustrated in Fig.3. In Fig.3.A, a characteristic bandpass filtered time series for each of the studying frequency band is estimated from each ROI. Here, instead of vectorising the upper triangular of an undirected FCG, we used the statistical and topological filtering FCG on its inherent format which is a 2D tensor. In the case of dynamic networks, the dimension is a 3D tensor where the 3^rd^ dimension is the time. Fig.3B illustrates a few snapshots of the dFCG for δ frequency of the first subject. At the next level, we estimated the laplacian matrix of each quasi-static FCG. Given a FCG, the laplacian matrix is given by:

L = D - A, (16)

where *D* is the degree matrix and A is the *FCG*.

Fig.3C demonstrates the laplacian matrix of the FCG in Fig.3B. In the main diagonal, the degree of each node is tabulated. Afterward, we applied an eigenanalysis for each of these laplacian matrixes and the eigenvalues of this procedure describes the synchronizability of the original FCG. Fig.3d illustrates for the 1^st^ minute the eigenvalues for each quasi-static FCG. One can easily detect the abrupt transition between the brain states. Here, neural-gas algorithm was applied on the 2D matrix presented in Fig.3D after first concatenated across subjects independently for each frequency band and scan session. The main scope of this codebook learning algorithm is to define FCμstates.

By estimating the reconstruction error E between the original 3D graphs and the one reconstructed via the k FCμstates assigned to each snapshot of the DFCG for each predefined threshold, we can detect the optimal threshold T for each case. In this work, the criterion of the reconstruction error E was set less than 4%. Practically for all the frequency bands and in both connectivity estimators, the reconstruction error E was less than 2%. The selected threshold was detected based on the plateau by plotting of reconstruction error E versus the threshold T.

In this way, the richness of information contained in the dynamic connectivity patterns is represented, by a partition matrix U, with elements u_ij_ indicating the assignment of input connectivity patterns to code vectors. Following the inverse procedure, we can rebuild a given time series from the k FCμstates, with a small reconstruction error E. The selection of parameter k reflects the trade-off between fidelity and compression level. As a consequence, the symbolic time series closely follows the underlying functional connectivity dynamics. The derived symbolic times series that keep the information of network FCμstates (nFCμstates) are called hereafter as STS^L-EIGEN^ (L:Laplacian – Eigen:Eigenalysis). Fig.3E tabulates the correlation of the eigenvalues between every pair of temporal segments while in Fig.3F, the matrix in Fig.3E was reordered such as the FCμstates to be revealed via the neural-gas algorithm. The network topology of the extracted FCμstates is illustrated in Fig.3G. From, Fig.3F, one can understand that the two FCμstates describe the DCFG of this subject.

An exemplar of prototypical FCμstates is illustrated in Fig.3.G. The outcome of this clustering procedure is also to extract a symbolic time series per subject, repeat scan and frequency that describes the transition of brain activity between the extracted brain states (FCμstates; Fig.3.D). The transition probability P for this example and for the two FCμstates is illustrated with a classical figure for Markovian Chain. The self-arrows refer to the percentage of sliding windows where the brain stays stable in a FCμstate without any transition while the directed arrow gives the percentage of transition from one FCμstate to the other. This symbolic time series can be seen as a Markovian chain where these switching between ‘quasi-static’ FCμstates can be modeled as a finite Markov chain (Dimitriadis et al., 2013a, b,2015a; O’-Neill et al.,2015; Vidaurre et al., 2016). One can clearly understand that human brain demonstrates a preferred transition from FCμstates^2^ to FCμstates^1^ (off-diagonal lines of the TP) compared to the opposite direction (Fig.3.H). The sketch of the markovian chain and the colored TP matrix can reveal the aforementioned trend of preferred direction FCμstates^2^ to FCμstates^1^.

From the symbolic timeseries, specific metrics tailored to the dynamic evolution of FCμstates were estimated (see next section) and their reliability was assessed via the correlation coefficient between scan session 1 and scan session 2. The whole approach was repeated independently for each frequency band and connectivity estimator by integrating subject and scan-based DFCG.

### 2.12 Characterization of time-varying connectivity

Once the integrated DFCG is formed and it is modelled via the combination of neural-gas and graph diffusion distance scheme (N-GAS^GDD^), relevant features can be extracted from the data based on the state-transition states. There features are called chronnectomics (chronos – Greek word for time and connectomics for network metrics) which are described in the following section.

### 2.13 Chronnectomics

The following chronnectomics (dynamic network metrics) will be estimated on the STS^L-EIGEN^ which expresses the fluctuation of the FCμstates.

#### 2.13.1 State Transition Rate

Based on the state transition vectors STS^L-EIGEN^ as demonstrated in Fig.3.A, we estimated the transition rate (TR) for every pair of states as followed:

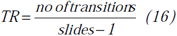

Where slides denote the number of temporal segments using the sliding window approach.

TR yields higher values for increased numbers of ‘jumps’ of the brain between the derived brain states over consecutive time windows. This approach leads to one feature per participant. Finally, we transformed the transition matrix (TM) into a probability matrix by dividing the TM by the sum of totally observed transitions over all possible pairwise states. TR was also estimated between every possible pair of states leading to an 8×8 TR matrix and extra 64 features per subject and condition.

#### 2.13.2 Occupancy Times of the nFCμstates

Complementary to the aforementioned chronnectomics, we estimated also the occupancy time (OC) of each FCμstates as the percentage of its occurrence across the experimental time. OC was estimated from STS^L-EIGEN^ as follows:

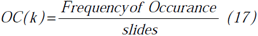

where k denotes the FCμstates.

### 2.14 Reliability of Static Network Metrics and Chronnectomics

The reliability of static node-wise network metrics and the chronnectomics was assessed with the correlation coefficient between forty values derived from scan session 1 and forty values from scan session 2 for each frequency band, condition and connectivity estimator (see Fig.1–3).

### 2.15 Optimizing the Width of the Time-Window and the Stepping Criterion

We optimized both the width of the time-window and the step criteria for the sliding-window approach based on the maximization of the reliability of TR. The reliability was estimated based on the correlation coefficient of the TR across the whole group between scan session 1 and 2. The whole procedure was followed independently for each brain rhythm. The settings for the width of temporal window and the step were defined as a percentage of the cycles of the studying frequencies: {from 1 up to 10 cycles with step equals to 0.5 cycle} for the width of the temporal window and {from 0.1 cycles to 2 cycles with step equals to 0.1 cycle} for the step.

To avoid overfitting of both TR and OT since, we used TR for both the optimization of the width of the temporal window and the stepping criterion, we used the optimized parameters in an external second repeat scan dataset for further evaluation.

## 3. Results

### 3.1 Tuning Parameters for Dynamic Functional Connectivity Analysis

The optimization of the temporal window and the stepping criterion for each brain rhythm reveals a nice trend for dynamic functional connectivity analysis. The width of the temporal window increased from δ to γ_2_ while the stepping criterion decreased in both connectivity estimators.

**Table 1.** Optimization of the width of temporal window and the stepping criterion per frequency band and for both connectivity estimators.

| | | $\delta$ | $\theta$ | $\alpha_1$ | $\alpha_2$ | $\beta_1$ | $\beta_2$ | $\gamma_1$ | $\gamma_2$ |
| --- | --- | --- | --- | --- | --- | --- | --- | --- | --- |
| iPLV | {time<br>window<br>,step} | {2,0.5} | {3,0.4} | {4,0.4} | {4,0.4} | {6,0.3} | {6.5,0.3} | {8,0.2} | {8.5,0.2} |
| CorEnv | {time<br>window<br>,step} | {2,0.5} | {3,0.4} | {5,0.4} | {5,0.4} | {7,0.3} | {7,0.3} | {8,0.3} | {9,0.2} |

### 3.2 Common Projection Space of Frequency-Dependent Static FCG

To demonstrate the (dis)similarities between sessions and subjects of the frequency-dependent static FCG, we constructed a distance matrix of dimensions 80 × 80 (subjects × sessions) using the graph diffusion distance metric. Then, we applied multidimensional scaling (MDS) to project the distance matrix into a common 2D feature space. Using a different coloured circle for each scanning session (blue for session 1 and red for session 2) and connecting both of them with a black line for each subject, we further enhanced the (dis)similarities of the static FCGs. Figure 4 – 5 illustrate these FCG-based projections for static FCG^Iplv^ and FCG^CorEnv^ correspondingly. In Fig.4G one can detect a few subjects with high reliable static FCG between the two scan sessions and also subject-specific network topologies that occupied an isolated subarea in the common projection FCG space. The stress index estimated via the MDS approach was low and the relationship of the 80 FCGs in the original 80×80 matrix is preserved in the projecte space.

**Figure 4.**
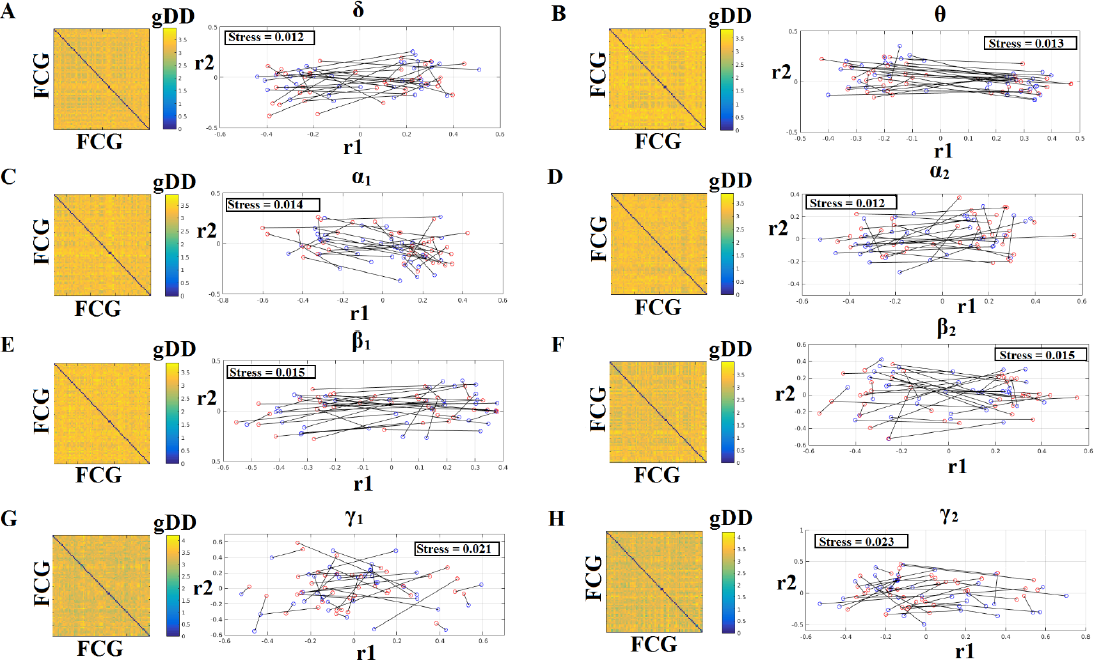
Muldimensional Scaling Projection of Frequency-Dependent Static Functional Connectivity Graphs (FCG^iPLV^) in a Common Feature Space. (A-H: δ - γ_2_) Each subplot illustrates the (dis)similarities of static FCGs across scanning sessions subjects. The 2D matrix demonstrates the (dis)similarities of the static FCGs across the subjects and repeat scans. Scanning sessions were coded with blue and red circles correspondingly and a blac connects the FCG of each subject between the two scanning sessions. With this representation on read out the similarity of a static FCG between two scanning sessions and participants.

Stress expresses the loss of information expressed in the projected Frequency-Dependent Functional Connectivity Graphs in 2D feature space from an original 80D space. The low stress v mean that the relationship of the 80 FCGs in the original 80×80 matrix is preserved in the projecte 2D space. R_1,2_ refer to the 2D projected space of the 80 FCGs.

**Figure 5.**
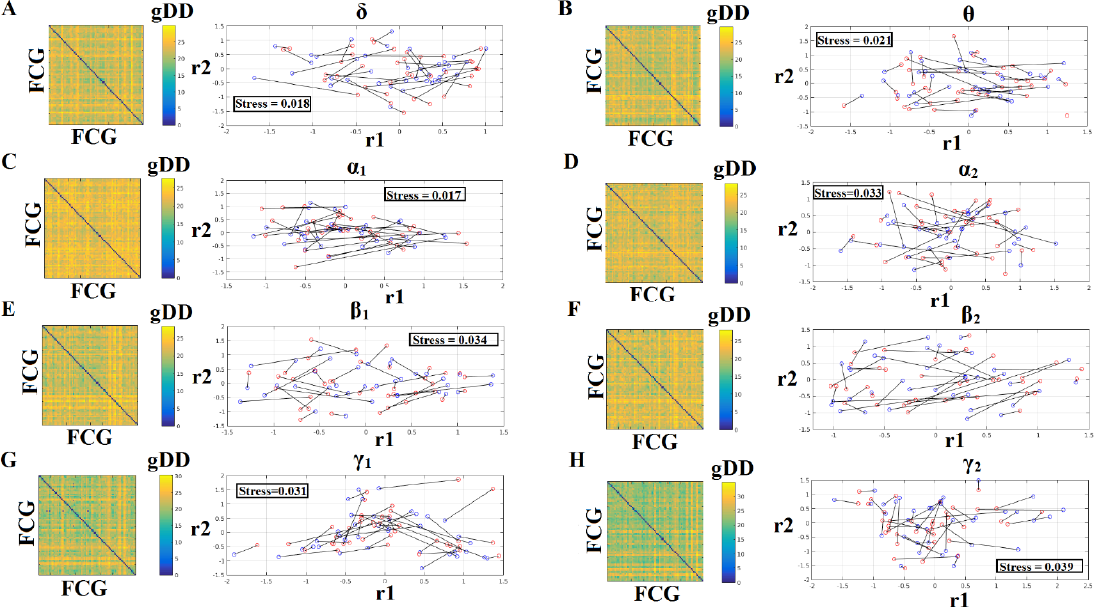
Muldimensional Scaling Projection of Frequency-Dependent Static Functional Connectivity Graphs (FCG^CorEnv^) in a Common Feature Space. (A-H: δ - γ_2_) Each subplot illustrates the (dis)similarities of static FCGs across scanning sessions subjects. The 2D matrix demonstrates the (dis)similarities of the static FCGs across the subjects and repeat scans. Scanning sessions were coded with blue and red circles correspondingly and a blac connects the FCG of each subject between the two scanning sessions. With this representation on read out the similarity of a static FCG between two scanning sessions and participants.

Stress expresses the loss of information expressed in the projected Frequency-Dependent Functional Connectivity Graphs in 2D feature space from an original 80D space. The low stress v mean that the relationship of the 80 FCGs in the original 80×80 matrix is preserved in the projecte space. R_1,2_ refer to the 2D projected space of the 80 FCGs.

### 3.3 Reliability of Static Network Metrics

Fig. 6 and 7 demonstrate the correlation coefficients for each node-wise network metric bet the two scanning sessions for every frequency-dependent static FCG. From the visual comparison of figures one can clearly reveal that the correlation values are higher for CorEnv compared to iPLV. App Wilcoxon Rank-Sum Test for every frequency and network metric between the 90 correlation value detected statistical significant differences in every case (p < 0.01, p’ < p/32, Bonferroni Corre However, the averaged correlation values did not reach high reliability (e.g. > 0.9) even for the CorE is obvious from the correlation plots that the reliability of node-wise static network metrics has spatial variability in both connectivity estimators.

**Figure 6.**
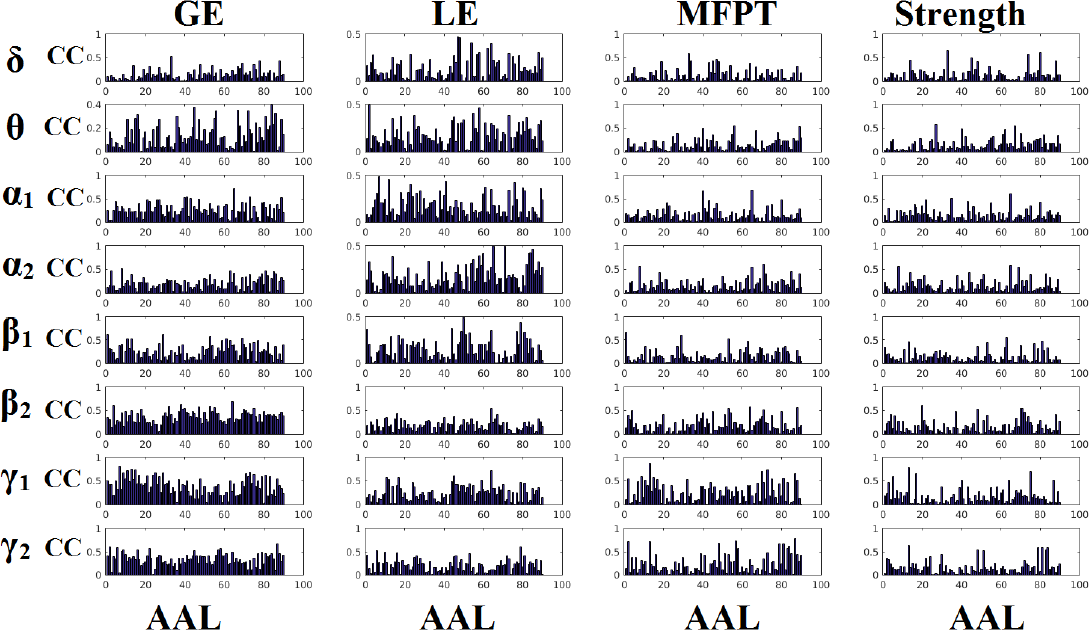
Reliability of node-wise network metrics derived from static brain networks with iPLV connectivity estimator. Each subplot demonstrates the correlation coefficient (CC) of each network metric at every studying frequency band of each brain area between the two scanning sessions. (CC: the correlation coefficient; AAL:Automated Anatomical Labeling)

**Figure 7.**
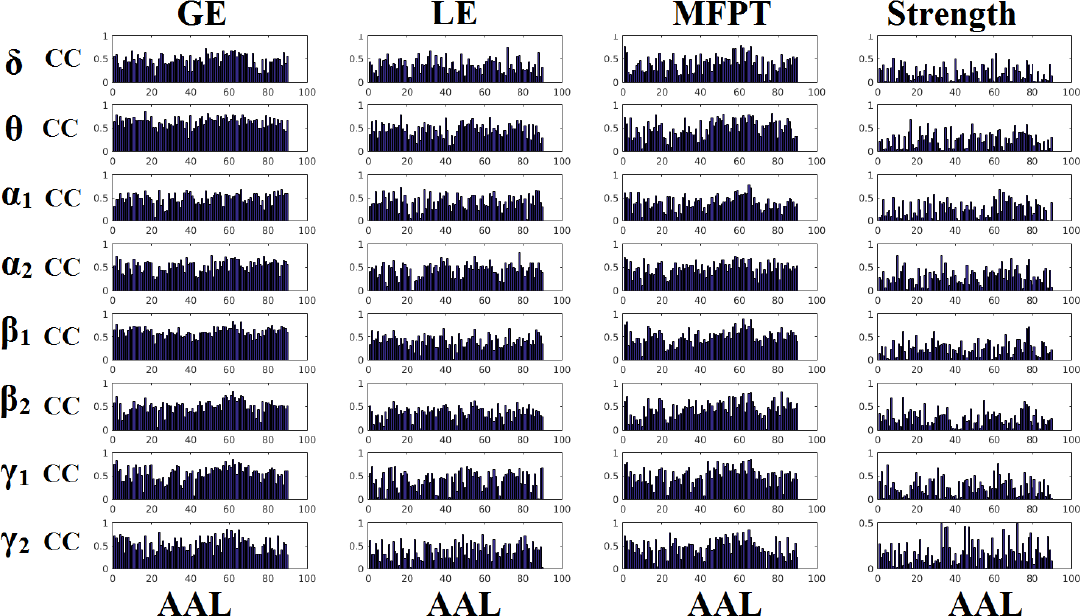
Reliability of node-wise network metrics derived from static brain networks with CorEnv connectivity estimator. Each subplot demonstrates the correlation coefficient (CC) of each network metric at every studying frequency band of each brain area between the two scanning sessions. (CC: the correlation coefficient; AAL:Automated Anatomical Labeling)

### 3.4 Frequency-Dependent FCGμstates and Reliability Chronnectomics for iPLV

Our analysis of DFCG based on iPLV revealed two FCGμstates^iPLV^ for each frequency band. The topology of these frequency-dependent FCGμstates^iPLV^ is illustrated in Fig.8. We integrated the nodes into five well-known brain networks: default-mode (DMN), fronto-parietal (FPN), occipital (O), sensorimotor (SM) and cingulo-opercular (CO). The mapping between the 90 ROIs of AAL and the five brain networks can be retrieved from section 3 in supp.material. One can clearly detect that the functional coupling between the default mode network and the cingulo-opercular dominates the coupling strength across the frequency bands and FCGμstates with less pronounced effect in both γ bands. Complementary, the coupling strength between and within the networks is diminished after α_2_ frequency. This behaviour can be interpreted as a reduction of the connections up to the defined threshold following the increment of the frequency. The two FCGμstates^iPLV^ showed also a different distribution of strength globally and locally.

**Fig. 8.**
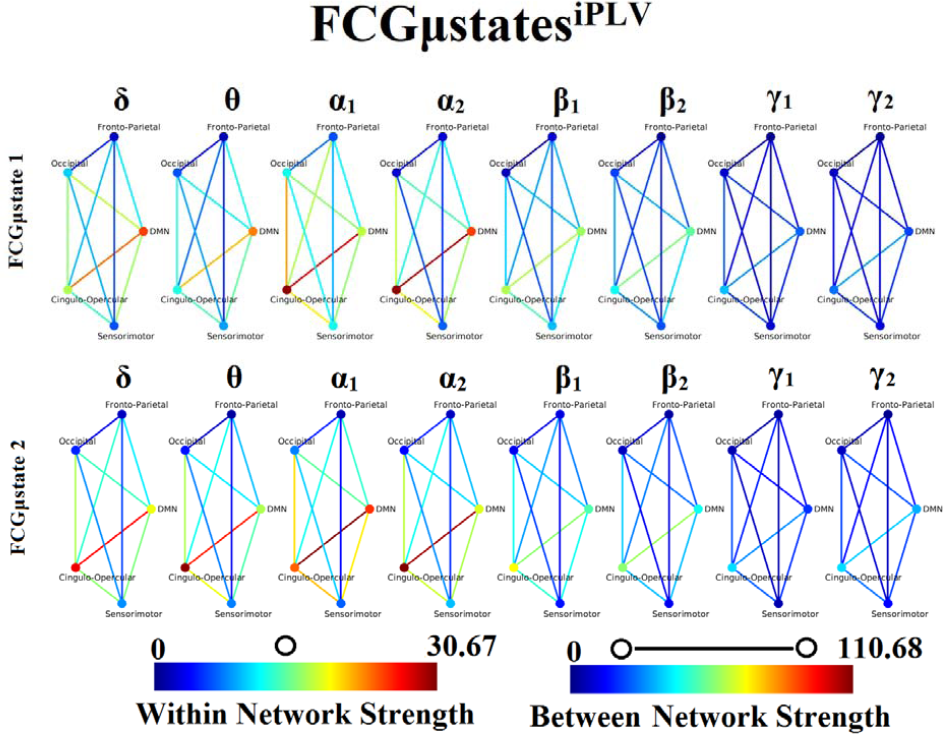
Frequency-dependent FCGμstates^iPLV^. Network topologies of the FCGμstates^iPLV^ for each of the studying frequency band. To enhance the visualization and contrast of FCGμstates across frequency bands, we adopted a netw wise representation instead of plotting the brain network in a 90 nodes layout. The 90 ROIs of the template were assigned to each of the five networks: default-mode (DMN), fronto-parietal (FPN), occ (O), sensorimotor (SM) and cingulo-opercular (CO). The color of each node denotes total streng within network connections while the color of each line the total strength of between net connections. Both strength values were normalized across both frequencies and FCGμstates.

Both types of chronnectomics, transition rates (TR) (Fig.9) and occupancy times (OC) (Fig.10) demonstrated high reliability (Corr > 0.9, p < 10^−7^) across frequency bands. Similar results, we obtained also for the second external dataset (see section 2 in sup.material).

**Fig. 9.**
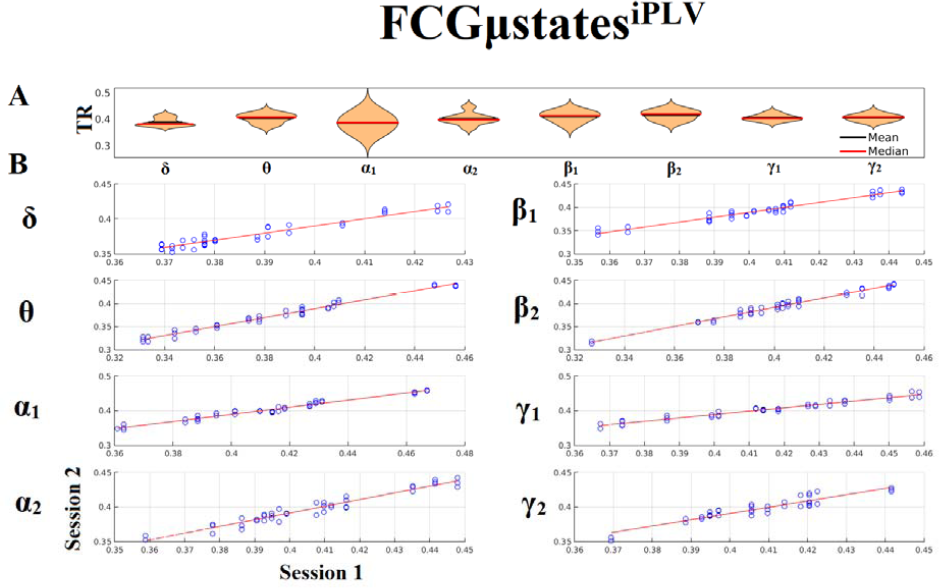
Reliability of Transition Rates (TR) based on FCμstates^iPLV^ across frequency bands. A. Mean and median values of TR across subjects and scan sessions for each frequency band B. Scatter-plot of subject-specific TR for both sessions with the corresponding fitted line for each frequency band. All the correlations were Corr.> 0.9 (p < 10^−7^) Each blue circle corresponds to a participant.

**Fig. 10.**
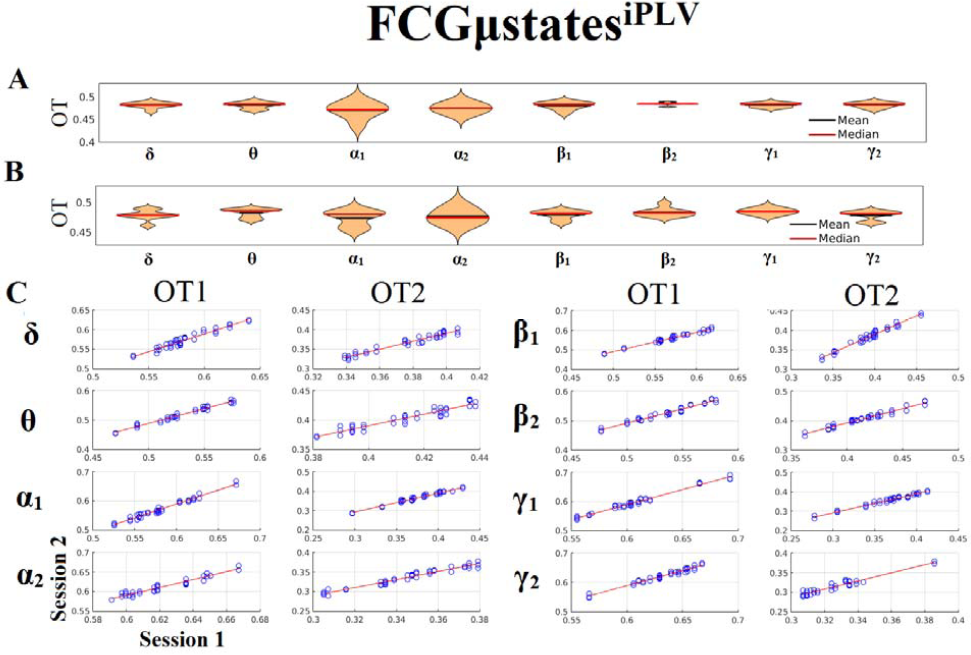
Reliability of Occupancy Time (OT) based on on FCμstates^iPLV^ across frequency bands. a. Mean and median values of OT across subjects and scan sessions for FCμstates^1^ and for ea frequency band b. Mean and median values of OT across subjects and scan sessions for FCμstates^2^ and for ea frequency band c. Scatter-plot of subject-specific OT for both sessions with the corresponding fitted line for e frequency band. All the correlations were Corr.> 0.9 (p < 10^−7^). Each blue circle corresponds to a participant.

### 3.5 Frequency-Dependent FCGμstates and Reliability Chronnectomics for CorEnv

Our analysis of DFCG based on the correlation of the envelope connectivity estimators revealed two FCGμstates^CorEnv^ for each frequency band. The topology of these frequency-dependent FCGμstates^CorEnv^ is illustrated in Fig.11. default-mode (DMN), fronto-parietal (FPN), occipital (O), sensorimotor (SM) and cingulo-opercular (CO). The mapping between the 90 ROIs of AAL and the five brain networks can be retrieved from section 3 in supp.material. One can clearly detect that the functional coupling between the default mode network and the cingulo-opercular dominates the coupling strength across the frequency bands and FCGμstates. Complementary, the coupling strength between and within the networks is diminished after α_2_ frequency as it was observed for FCGμstates^iPLV^. Complementarily, the network topologies of FCGμstates^CorEnv^ between low and high frequencies basad on the strength coupling are more common than the FCGμstates^iPLV^. This common substrate across the FCGμstates^CorEnv^ is consistent with the general notion that correlation of the envelope is more stacked to the structural connectome compared to the phase-based connectivity patterns which demonstrate higher degrees of freedom (Engel et al., 2013; compare Fig.8 with Fig.11).

**Fig. 11.**
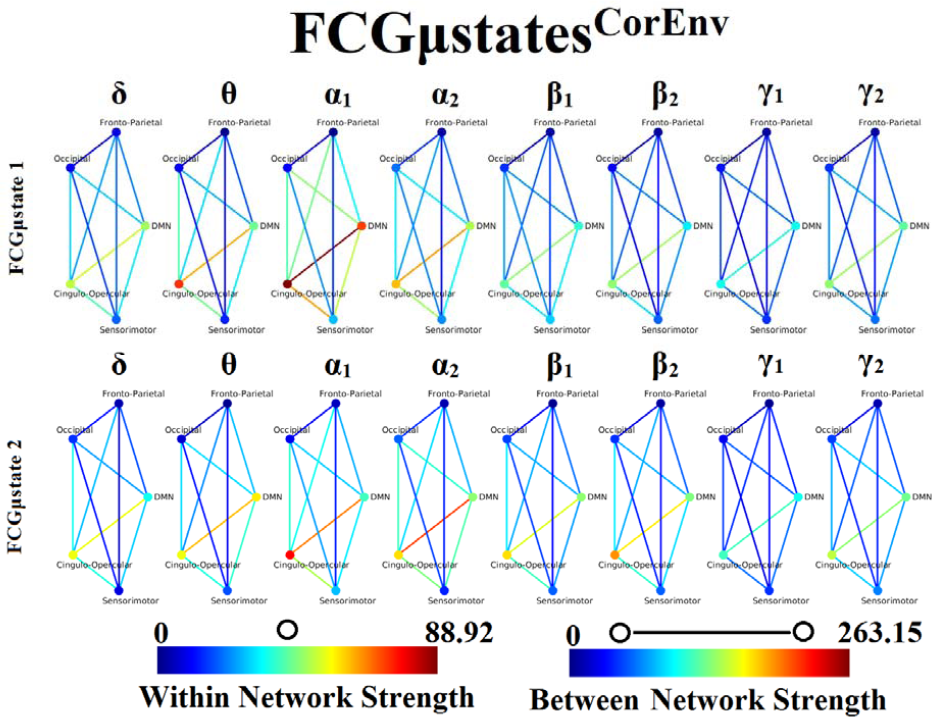
Frequency-dependent FCGμstates^CorEnv^. Network topologies of the FCGμstates^CorEnv^ for each of the studying frequency band. To enhance the visualization and contrast of FCGμstates across frequency bands, we adopted a netw wise representation instead of plotting the brain network in a 90 nodes layout. The 90 ROIs of the template were assigned to each of the five networks: default-mode (DMN), fronto-parietal (FPN), occ (O), sensorimotor (SM) and cingulo-opercular (CO). The color of each node denotes total streng within network connections while the color of each line the total strength of between net connections. 7Both strength values were normalized across both frequencies and FCGμstates.

Only transition rates (TR) showed high reliability for CorEnv (Cor.> 0.8, p < 10^−4^) in all the frequency bands with the only exception of β_1_ (Fig.12). Occupancy times (OT) showed low reliability across the frequency bands (p > 0.4)(Fig.13). TR of FCGμstates^CorEnv^ increased with the increment of frequency reaching a plateau in β_1_. In contrast, TR of FCGμstates^iPLV^ did not show such a frequency-dependent behavior. Similar results, we obtained also for the second external dataset (see section 2 in sup.material).

**Fig. 12.**
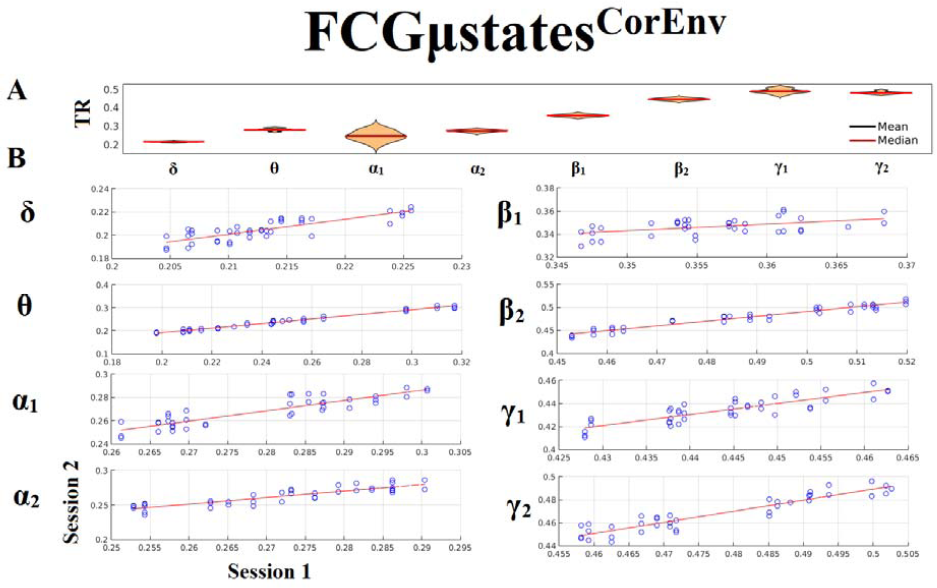
Reliability of Transition Rates (TR) based on FCGμstates^CorEnv^ across frequency bands. a. Mean and median values of TR across subjects and scan sessions for each frequency band b. Scatter-plot of subject-specific TR for both sessions with the corresponding fitted line for each frequency band. All the correlations were with the exception of β_1_ were Corr.> 0.8 (p < 10^−4^). c. Each blue circle corresponds to a participant.

**Fig. 13.**
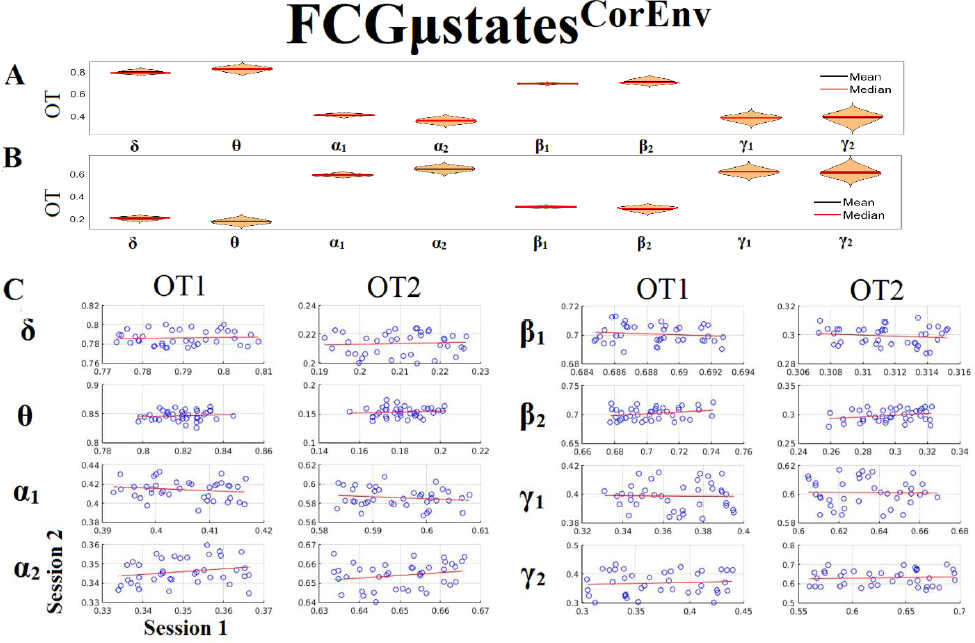
Reliability of Occupancy Time (OT) based on on FCGμstates^CorEnv^ across frequency bands. a. Mean and median values of OT across subjects and scan sessions for FCμstates^1^ and for each frequency band b. Mean and median values of OT across subjects and scan sessions for FCμstates^2^ and for each frequency band c. Scatter-plot of subject-specific OT for both sessions with the corresponding fitted line for each frequency band. All the correlations were weak and non-significant(p > 0.4). d. Each blue circle corresponds to a participant.

## 4. Discussion

In the present study, we assessed the reliability of both static and dynamic functional connectivity network descriptors using resting-state MEG data from 40 subjects with repeat scan sessions. This is the first time that the reliability of chronnectomics, at least for the MEG modality, has been taken into account. Source time series were first beamformed independently for each frequency band (Brookes et al., 2011b; Hillebrand et al., 2005; Schoffelen and Gross, 2009), and then representative voxel time series based on the AAL atlas were extracted using a novel linear interpolation analysis. This procedure produces informative timeseries with a characteristic carrier frequency compared to the noisy time series derived by PCA or by selecting the voxel time series within a ROI that encapsulates the maximum power. Then, both static and dynamic frequency-dependent functional connectivity graphs were computed for each subject and scan session using the imaginary part of phase locking value (iPLV) and the correlation of the amplitude envelope (CorEnv). Both static and dynamic FCG (SFCG-DFCG) were filtered both statistically (surrogates) and topologically (OMST; Dimitriadis et al., 2017a, b).

Here, we adopted a data-driven pipeline of how to estimate both static and dynamic FCG statistically and topologically filtered using an algorithm previously applied to EEG recordings. We explored the reliability of both static network metrics and chronnectomics (dynamic network metrics) by employing two representative connectivity estimators for the construction of static and dynamic brain networks. Using this pipeline, prototypical FCμstates were derived which were highly reproducible across subjects and scan sessions in both connectivity estimators and in all frequencies. The reliability of node-wise static network metrics based on four network metrics was low and spatially variable with both connectivity estimators while the CorEnv demonstrates higher ICC values compared to iPLV. The reliability of chronnectomics (TR, OT) for iPLV was high while for CorEnv the reliability of only the TR reaches high acceptable levels. Our results were reproduced also in a second external dataset (see supp. material). Our study strongly encouraging the use of DFCG with neuromagnetic recordings that takes the advantage of the nature of MEG modality, its high temporal resolution.

To our knowledge, this is the very first study that explored the reliability of both static and dynamic FCG and the related network metrics and chronnectomics, respectively in neuromagnetic source space at. In static FCG, node-wise network metrics demonstrated poor reliability for iPLV and poor to medium for CorEnv. The node-wise reliability was highly spatial variable and static FCG have also demonstrated low repeatability in both connectivity estimators and especially in CorEnv. In contrast, prototypical FCμstates were high reproducible across subjects and scan sessions in both connectivity estimators and in all frequencies supporting by the low reconstruction error (< 2%) of our brain network learning algorithm. Complementary, the reliability of chronnectomics (TR, OT) for iPLV was high while for CorEnv the reliability of only the TR reaches high acceptable levels. These results strongly encourages the neuroscientists to adopt the notion of DFCG with neuromagnetic recordings that takes the advantage of its high temporal resolution.

Two main studies explored the reliability of static FCG on the source level using MEG-beamformed resting-state connectivity analysis. Garces et al., (2016) studied the reliability of resting-state networks using four connectivity estimators: phase-locking value (PLV), phase lag index (PLI), direct envelope correlation (d-ecor), and envelope correlation with leakage correction (lc-ecor). They adopted intra-class correlation coefficient (ICC) and Kendall’s W for assessing within and between-subjects agreement respectively. Higher test-retest reliability was found for PLV from θ to γ, and for lc-ecor and d-ecor in ². They commented that high ICC in PLV and d-ecor could be artifactual due to volume conduction effects. Colclough et al., (2016) investigated the reliability of static FCG at resting-state using beamformed source static connectivity analysis. They reported high reliability mostly for the partial correlation analysis and the correlation of the envelope among 12 connectivity estimators. Two more studies, Deuker et al., (2009) estimated the reliability of resting-state network metrics derived from MEG in sensor space using mutual information. They obtained high ICC for clustering, global efficiency and strength at a network level. Jin et al.,(2011) found medium ICC for nodal global efficiency, nodal degree and betweeness centrality in α and β bands.

Our results revealed that nodal network metrics derived from static FCG are less reproducible then their dynamic counterparts. In contrast, chronnectomics are highly reproducible with both adopted connectivity estimators. These results complemented with the results presented in (Colclough et al., 2016) where they adopted multiple connectivity estimators for the construction of static brain networks on the source level using MEG-beamformed resting-state activity. Colclough et al., (2016) showed that the static full-weighted FCG are high repeatable within the group-level mostly for the correlation of the envelope adopting a split-half strategy on a dataset with single scans. Here, we accessed the reliability of any network metric using a two scan session design per subject. We should state here that edge-weights are significant for the construction of network topology and the reliability of connectomic biomarkers (Dimitriadis et al., 2018).

One of the key findings of our analysis are the frequency-dependent FCμstates for each connectivity estimator. Figures 8 and 11 illustrate the strength of the coupling within and between brain networks for the prototypical FCμstates at every frequency band. It is obvious that the highest strength within a network is observed within the DMN in both connectivity estimators. The strength between the brain networks is mainly distributed between DMN and the rest of the networks demonstrating the highest value till α_2_ and dropped from β_1_ to γ_2_ (Figs.8,11). DMN reignited a high interest the last years for the description of intrinsic ongoing activity in studies of the human brain in health and disease (Raichle et al.,2015). Disruptions of functional connections within the DMN and between DMN and the rest of brain networks has been linked to various neurological and neuropsychiatric disorders (Mohan et al., 2016). Studies in healthy aging and Alzheimer’s disease have revealed the significant role of DMN (Mevel et al., 2011).

Flexible hub theory based on clustering analysis of functional networks gave an explanation of how temporal functional modes exist where one neural region may be switched from a certain network at one time to a different network at another time (Smith et al., 2012). It remains still unclear how the different brain networks are connected together during spontaneous and task-related activity. Dosenbach et al. (2008) proposed that the FPN may serve to initiate and adjust cognitive control, whereas another control-type network, the CO network (CON), provides stable set-maintenance. Cole and colleagues (Cole et al., 2013) helped to untangle the flexible role of the FPN, many questions remain regarding the interaction between the FPN and the CON and also with other networks such as the DMN, SM and O. In the present study, we characterized the dynamic relationships of the brain networks across time at resting-state in various frequency bands and using representative connectivity estimators. We found that these functional patterns are high reproducible which will help multi groups worldwide to explore these interactions and build reproducible connectomic biomarkers in various diseases and disorders. Understanding the neural basis of intrinsic activity, cognition and structure–function relationships, will further enhance the prognostic/diagnostic abilities in clinical populations.

The interactions of large-scale brain networks at resting-state and during tasks is characterized by the studying frequency. Frequency-specific functional interactions between large-scale brain networks may be individual fingerprints of the idle activity and cognition (Siegel et al., 2012). It will be interesting in the future to explore how the FCμstates from a dynamic integrated functional connectivity graph (Dimitriadis et al., 2017c) that incorporates different intrinsic coupling modes both intra and cross-frequency coupling can be used for brain fingerprinting (Engel et al., 2013).

It is critically important to take advantage of new imaging modalities to untangle the mechanisms that produce circuit dysfunctions in many brain diseases and disorders. The development of biomarkers is very important and for that reason the proposed experimental paradigm and analytics of the meta-data derived from the analysis of human brain activity must be highly reliable and reproducible. Magnetoencephalography (MEG) allows us to measure neuronal events noninvasively with millisecond resolution and recent advanced methods opens new avenues of exploring and answering fundamental key research questions tailored to each brain disease/disorder. MEG can become a pioneering clinical research tool for mental disorders (Bowyer et al., 2015; Grent-T-Jong et al., 2016) (Uhlhaas et al., 2017), Alzheimer’s disease (Lopez et al., 2014,2017; Koelewijn et al., 2017), dyslexia (Dimitridis et al., 2013b,2016b), traumatic brain injury (Dimitriadis et al., 2015c, Antonakakis et al., 2016, Antonakakis et al., 2017), multiple sclerosis (Tewarie et al., 2015), and other brain diseases. To establish MEG-based biomarkers that can be used for daily clinical practise and clinical evaluation, their reproducibility should be further explored. Complementary, the transition rate and also the occupancy times could be personalized biomarkers of a subject’s resting-state condition where more task-related FCμstates and the related markers derived from them could build a subject specific database for longitudinal studies. Transition rates could be also correlated with IQ scores and also with behavioural performance during execution of cognitive tasks.

In the present study, we proposed a data-driven analytic pathway to assess the reliability of connectomics using MEG-beamformed connectivity analysis. Our results clearly support the notion of dynamic functional connectivity on the source level, the derived prototypical FCμstates and the related chronnectomics. Last years, many studies explored the dynamic functional connectivity graphs in many modalities (EEG/MEG/fMRI) and in both resting-state and during tasks (Dimitriadis et al., 2015b; Mylonas et al., 2015; Toppi et al., 2015; Yang and Lin, 2015; Calhoun and Adali, 2016, for reviews see Calhoun et al., 2014). This is the very first study according to authors’ knowledge that the reliability of chronnectomics was explored. The outcome of this study opens new avenues in the exploration of human brain dynamics with MEG-beamformed source activity and under the notion of dynamic functional connectivity.

We addressed the key question of the readiness of neuromagnetic-based based functional connectomics to lead to clinically meaningful biomarker identification through the reliability approach that offers a repeat scan study in healthy controls. It is more than significant to customize stable approaches for analysing neuromagnetic recordings and present reproducible brain connectomics across scans in healthy control populations without sacrificing the individual characteristics that can be used for personalized intervention neuroscience (Gratton et al., 2018). It is highly recommend to access the reliability of any metric derived from any neuroimaging modality in a repeat scan protocol in healthy control population before applying it to a larger disease group where the cost of scanning is too high (diffusion MRI: Dimitriadis et al., 2017d). Additionally, we will expand this analysis in future efforts to identify disease status alone including clinical variables related to genetic risk (Lancaster et al., 2018), expected treatment response and prognosis.

## 5. Conclusions

In conclusion, we provided the first source-space test-retest reliability of dynamic functional connectivity of neuromagnetic recordings at resting-state. We computed both static and dynamic functional connectivity based on 90 ROIs according to AAL templated and using two connectivity estimators, the iPLV and the CorEnv. Nodal network metrics were unreliable in both connectivity estimators but with higher reliability demonstrated for CorEnv. Moreover, their reliability demonstrates highly spatial variability. Static FCG were also unreliable and especially for CorEnv. In contrast, prototypical FCμstates were reliable in both connectivity estimators and across frequency bands. The derived chronnectomics (TR, OT) were highly reproducible for iPLV while only TR was reliable for CorEnv within acceptable levels. Our results strongly encourages future studies with main scope to explore resting-state networks in both healthy control and disease populations to apply a data-driven dynamic functional connectivity analysis using MEG-beamformed source reconstructed brain activity.

## Acknowledgements

SID and DL were supported by MRC grant MR/K004360/1 (Behavioural and Neurophysiological Effects of Schizophrenia Risk Genes: A Multi-locus, Pathway Based Approach). SID is also supported by a MARIE-CURIE COFUND EU-UK Research Fellowship. BR and the CUBRIC MEG lab are supported by an MRC UK MEG Partnership Grant, MR/K005464/1 and an MRC Doctoral Training Grant, MR/K501086/1. We would like to acknowledge RCUK of Cardiff University and Wellcome Trust for covering the publication fee.

## Author Contributions

Conception of the research analysis: SD; Methods and design: SD; Data analysis (SD); Drafting the manuscript: SD; Data Acquisition:(BR);Critical revision of the manuscript: (BR, DL, KS); Every author read and approved the final version of the manuscript.

